# “It’s not just about the science”. The impact of undergraduate research projects and COVID-19 on graduate attributes and employability

**DOI:** 10.64898/2026.02.13.705786

**Authors:** Maria Kyriazi, Jessica F. Jung, Simon Wilkinson, Alistair K. Brown, Katerina Panti, Vanessa L. Armstrong

## Abstract

Over the past two decades, Higher Education Institutions have increasingly prioritised transferrable skills to enhance graduate employability. Graduate Attributes (GAs) now act as key indicators of student competencies for both learners and employers. Final-year research projects, typically high in credit value, represent capstone experiences that promote subject expertise and GA development through research, written work, and oral presentations.

This study analyses pre- and post-project survey data from RQF Level 6 biomedical and biomolecular science students at a Russell Group University over four years (2019–2023). Most projects were laboratory-based, though the 2020–2021 cohort completed theirs remotely due to COVID-19.

Students reflected on expectations and experiences of GA development, subject knowledge, and employability. Initial responses revealed anxiety and uncertainty, particularly among the 2020–2021 cohort, but most anticipated gains in skills and employability. Post-project feedback confirmed this, identifying critical thinking, confidence, resilience, collaboration, and future focus as key outcomes. Digital capability was notably strengthened, especially during remote delivery.

The findings emphasise the importance of a shared understanding of GAs in bioscience education and the value of embedding structured reflection and preparatory support to help students recognise and articulate their evolving skills.

## Introduction

Graduate attributes (GAs) represent the essential transferable skills that students are expected to develop throughout their studies, which align with the values and strategic vision of Higher Education Institutions (HEIs) (Barrie, 2007; Bridgstock, 2009; Hill et al., 2016; Kotsiou et al., 2022; University, 2018). These are often termed as *soft skills* and encompass all the higher-level competencies that support students academically during their studies, but also professionally post-graduation. It is widely acknowledged that GAs do not solely enhance employability, but they also foster citizenship and promote teamwork and communication (Jakubik et al., 2023; Oliver & Jorre de St Jorre, 2018). External pressures from governments and employer stake holder groups have driven universities to adopt GAs with an expectation to improve graduate skill sets to enhance (Clarke, 2018). Graduate outcomes also form a key element of university performance indicators, including the Teaching Excellence Framework, which measure the impact on recruitment and the sustainability of institution and course programmes. In this context, modern graduates are expected to evolve diverse skill sets and capabilities that will facilitate their navigation to the complex, dynamic, and unpredictable workforce (Clarke, 2018). Understanding the benefits of their disciplinary perspectives, workplace inequalities, and retaining identity are challenges that graduates face. Therefore, successful GA acquisition allows them to respond to these challenges and thrive in ever-evolving environments.

Despite the importance of GAs, there is no defined list or terminology across the sector. Nevertheless, critical thinking, problem solving, digital literacy, preparation for lifelong learning, and resilience feature in most institutional models. The recently developed triple helix model (subject, object and world being core) lists 17 essential skills for graduate development (Ehlers, 2024). However, the absence of standardised terminology combined with the need for HEIs to tailor the framework to their institution, can lead to inconsistencies and confusing nomenclature. To address this issue, Advance HE has introduced a framework which incorporates behaviours, qualities and values; professional identity and networks; emotional intelligence, agency and self-awareness; confidence, adaptability and situational knowledge as key attributes that should be embedded within degree programmes (HE, 2024). In some cases, GAs have been replaced with models of graduate capital, particularly in Australia, which include human and social capital, individual attributes and behaviours, and perceived employability (Clarke, 2018).

GA frameworks need to be agile and responsive to the rapidly changing demands of the global markets. Adaptability and flexibility are critical, as evident by the impact of COVID-19 on education and employment, which highlighted the need for graduate preparation during times of uncertainty and rapid change (Gow & McDonald, 2000; Tomlinson et al., 2023). Since COVID-19, the nature of work has changed, characterised by accelerated technological development and an agile workplace. According to the World Economic Forum, this shift prioritises, technological adaptability, and predicted changes in workers’ skillsets with a focus on continuous learning, upskilling and reskilling to enable businesses to adapt and manage future skill requirements (Forum, 2025). Large data sets, cloud computing, and artificial intelligence are recognised as essential skills with analytical skills and creative thinking, identified as key attributes required. As many institutions reconsider their offer in the current financial situation, the purpose, value and role of Higher Education is questioned alongside this, emphasis in the articulation and drawing out where these skills and attributes are already identified in the curriculum is of key importance (Daubney, 2022). This reassessment, along with the fast-evolving workplace, has brought the reviewing of GAs to the forefront (Donald, 2025; School of Biomedical and Nutritional Sciences, 2025). In parallel, a skills gap reported by employers has reinforced the need for universities to ensure that GAs are clearly identified, understood by students, and then subsequently demonstrated to employers [16-18]. According to the 2022 QS report, globally, employers deem communication, teamwork, problem-solving and flexibility most highly (Symonds, 2022).

Newcastle University developed its Graduate Framework in 2018 following consultation with key stakeholders across the sector (University, 2018). Ten defined attributes were identified with global, culturally aware, reflective and self-aware overarching this framework, study (Figure 1).

**Figure 1:**
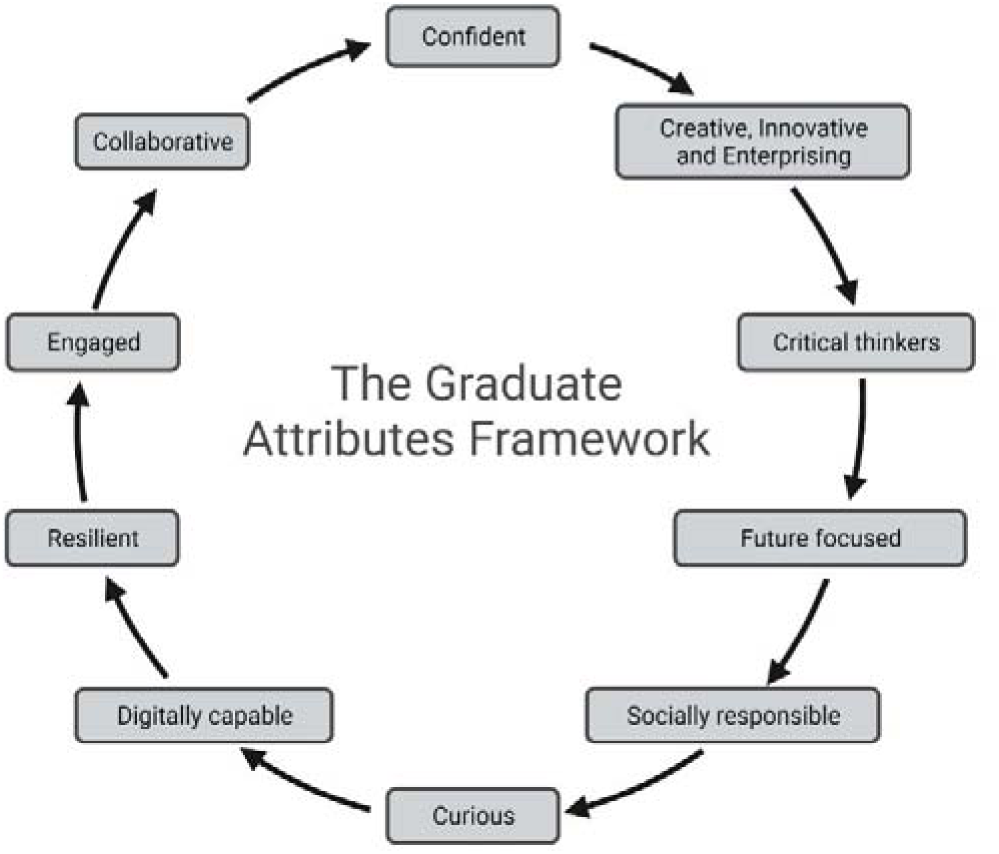
Newcastle University Graduate Attributes Framework (University, 2018).

Importantly these GAs are embedded institution-wide, rather than being discipline-specific, representing a core outcome of Higher Education that every graduate is expected to possess regardless of their subject area (Barrie, 2007). The Quality Assurance Agency (QAA) benchmark for biosciences, also identifies problem identification and solving, innovation, creativity, communication and practical action as skills that should be developed by students, enabling the transition into successful graduates (Membership, 2023). Therefore, given the extensive number of varying attributes and the lack of clear consensus across the sector, albeit with distinct similarities and cross over between these definitions, the GA offer needs clear articulation. Defining clearly which GAs should be emphasised and classifying them in a subject and institutional context is essential for ensuring stakeholder alignment and active engagement of the GAs in the syllabus (Oliver & Jorre de St Jorre, 2018).

GAs can be developed through formal curriculum design and extracurricular opportunities, such as sport, volunteering, work placements and part-time jobs (Buckley & Lee, 2021; Christison, 2013; Dickinson et al., 2021; Fakhretdinova et al., 2021). Recent research by the Office for Students, confirms that students value acquiring knowledge, as well as, developing skills that will enhance their career prospects (Students, 2025), and recognise the significance of developing transferable skills across their studies (Haigh & Kilmartin, 1999).

Extended research projects, or dissertations, are usually completed during the final year of undergraduate studies. During this period, students are encouraged to actively participate in research questions which require in-depth analysis of a problem. Research projects enable students to develop skills including creativity, independence and often teamwork, problem solving, and resilience, which are pivotal for future employability as well as actively implementing degree-specific knowledge (Healey et al., 2013; Lopatto, 2007; Seifan et al., 2022). Although originating in the USA, the Boyer Commission’s 1998 report *“Educating Undergraduates in the Research University”* outlined ten key recommendations for higher education institutions, notably advocating that “research-based learning” should become the norm, - a goal effectively realised through the introduction of undergraduate research projects (Undergraduates., 1998).

Most university institutions worldwide provide the opportunity to undertake an undergraduate research project, with slight variations across different countries. In the UK, dissertations are part of a traditional honours project timetabled to span the final year or one semester of the academic year. During this time, students work autonomously under the supervision of an academic staff member to address a research question and document the results of their research with a formal report (Rowley & Slack, 2004). The dissertation usually comprises a written thesis of approximately 5,000 to 12,000 words, alongside a presentation, and generally accounts for 30-60 credits of the total 120 credits required for Regulated Qualifications Framework (RQF) Level 6. It also gives students the opportunity for novel assessment exposure, GAs development, as listed above, and cultivation of subject-specific skills, such as literature review and research synthesis, bringing together topics previously covered in the course. The advantages of an undergraduate research experience are well established in the literature. These include numerous learning gains, improved academic performance (Fechheimer et al., 2011), higher retention levels and science pathways for minority students (Lopatto, 2007) and support for progression to further study (Hathaway et al., 2002; Lopatto, 2004). The importance of educators highlighting the innate employability value of what is already in the curriculum is emphasised by Daubney (Daubney, 2022).

Given that biomedical and biomolecular graduates pursue a variety of careers and further study opportunities post-graduation, some GAs may be more relevant than others. Data from the School indicates that further study is the most popular route for graduates in this discipline, with continuation into a master’s research course being the most popular option.

The Royal Society of Biology defines a capstone experience as a minimum of 30 credits and will be; “a. An extended piece of enquiry-based work, relevant to the degree, with a justified approach that effectively communicates the outcome, b. Underpinned by a range of relevant sources, and will show recognition of health and safety, environmental and ethical considerations, c. Contextualised, and show recognition of the provisional nature of knowledge, building to an appropriate conclusion, d. Based on the process of critical thinking, synthesis, reflection and evaluation”(Biology). Within Biomedical and Biomolecular Sciences at Newcastle University, students undertake a 40-credit capstone research project during semester 2 of Year 3 (RQF Level 6, January to April). This equates to 22% of marks contribution towards the final degree. For 10 weeks, students immerse themselves in their research project, complete a 5,000-word report (75% of module mark), deliver a 10-minute oral presentation (15%) and receive a professionalism assessment from their supervisor (10%) (School of Biomedical and Nutritional Sciences, 2025). For RQF Level 6 MSci (integrated masters) students, the structure remains the same as that for BSc students but has a different code. Over the period of this study, apart from the academic year 2020-2021 due to COVID-19 restrictions, just over three quarters of projects were laboratory-based, involving hands-on, ‘wet’ research. The first lockdown occurred after the completion of projects in March 2020 (2019-20 cohort), consequently only the 2020-21 cohort had 100% ‘dry’ non-laboratory-based projects, which were conducted remotely due to the local restrictions. Project topics and titles are set by the Principal Investigator, based across the Faculty of Medical Sciences within one of the three research institutes. Non-lab projects may cover epidemiological and psychosocial aspects of health, anonymised patient data, genomics, proteomics, systems biology studies that incorporate bioinformatics, or education-based projects. Ensuring that all projects, whatever type, are hypothesis driven and involve the analysis and critical evaluation of hypothesis driven research, is key to achieving Royal Society of Biology accreditation which these 10 programmes have. (Biology).

Despite the extensive research available on GAs and frameworks in the literature, it remains unclear whether all stakeholders fully understand the meaning and relevance of these attributes, particularly across various disciplines (Rook & Sloan, 2021). There is also lack of literature focusing on skill acquisition through research projects, particularly within non-clinical life science courses.

### Study aims

This study sought to explore students’ perceptions of their upcoming high stake research project, with particular emphasis to GA development and its potential influence on students’ perceived employability. A key objective was to create a structured opportunity for students to reflect on the skills obtained during their final year and consider their next steps. The timing of this research project, directly before the Easter break with students returning briefly before undertaking their final exams means that it a particularly busy and stressful time for students and they may not have had time or space to reflect on the recent skills and attributes that they had developed. In addition, as discussed above, there is the question as to whether all stakeholders understand what is meant by the GAs and are all prospective graduates able to articulate how they have developed these competencies.

Finally, we examined the impact of COVID-19, the transition from in person to remote project delivery and we assessed its implications for skill development and student experience. Our findings will inform local practice, evidence which GAs are linked to research projects, impact on perceived employability, subject knowledge and ultimately graduate employability that can be developed and enhanced via the capstone research project experience. There is also scope across disciplines and institutes to explore the GA gains and value of capstone research projects from a skills and employability perspective.

## Materials and methods

Data were collected over four academic years, from 2019-20 to 2022-23, within the School of Biomedical, Nutritional and Sport Sciences at Newcastle University. Participants included RQF Level 6 and Level 7 students enrolled in bioscience and biomolecular degrees, in particular, Bachelor of Science (BSc) and Integrated Masters (MSci) courses in Biomedical Sciences, Biochemistry, Biomedical Genetics, Physiological Sciences and Pharmacology. The cohort size ranged from 250 to 350 students annually, and the survey was distributed to those students in Year 3 (RQF Level 6).

Two surveys were run each academic year, a pre-project survey in semester 1 and a post-project survey during semester 2, following the completion of all dissertation components. Except for the first survey iteration, conducted in person during a scheduled teaching session using Ombea software, all subsequent surveys were conducted online via SurveyMonkey and disseminated through university email and the Virtual Learning Environment (Canvas). Questionnaires consisted of ten closed and one open-ended question seeking information regarding student demographics and background, expectations, feelings, and perceptions about the upcoming research project (pre-project), as well as reported skills development (post-project). All questions were aligned with the Newcastle University Graduate Framework (University, 2018).

### Ethics and consent to participate

Ethical approval was obtained from the Faculty of Medical Sciences Ethics Committee at Newcastle University (reference: 2019-12487) prior to the study in 2019. Participants provided written consent, were informed that the survey completion was voluntary and had the right to withdraw from the study at any point. Rationale and full project details were provided at the start of online survey. No sensitive data that could be attributed to a student was collected. All survey data were analysed anonymously and no survey linking between the pre- and post-project survey was carried out.

Data were analysed using Microsoft Excel and all graphs were plotted using GraphPad Prism software version 8.3.1. Statistical analysis (non-parametric chi-squared tests) carried out using GraphPad Prism software version 8.3.1. The numerical count of responses was weighted to account for the different sample sizes of the responders across the four cohorts. A threshold of P=0.05 was set and P values <0.05, <0.01, <0.001 and <0.0001 (*, **, ***, ****) were considered statistically significant. Values >0.05 were deemed as non-significant.

## Results

### A) Cohort demographics

The demographic data for the student profiles and participation in the surveys fluctuated from 16.2-35.9% of the total students per cohorts (Table 1). The percentage of female and male responders ranged, with female participants accounting for 71-83% of student responders, whereas males for 11-26%.

**Table 1.** Student characteristics. Pre-project data are presented in black and post-project results are in blue. Please note that not all participants responded to all of the demographic questions resulting in small variabilities across demographics.

| Student demographics: | 2022-2023 | 2021-2022 | 2020-2021 | 2019-2020 |
| --- | --- | --- | --- | --- |
| Responses (% of cohort) | 72 (21.6%) | 77 (20.2%) | 70 (19.2%) | 122 (35.9%) |
|  | 54 (16.2%) | 76 (19.9%) | 71 (19.5%) | 82 (24.2%) |
| Female | 80.6% (n=58) | 83.1% (n=67) | 71.4% (n=18) | 71.31% (n=25) |
|  | 72.2% (n=39) | 88.2% (n=67) | 81.7% (n=58) | Data not collected |
| Male | 19.4% (n=14) | 11.7% (n=9) | 25.7% (n=18) | 20.5% (n=25) |
|  | 24.1% (n=13) | 11.8% (n=9) | 11.6% (n=12) | Data not collected |
| Binary/third gender | 0% | 0% | 0% | 0% |
|  | 0% | 2.6% (n=2) | 2.9% (n=2) | Data not collected |
| Prefer not to say | 0% | 2.6% (n=2) | 0% | 0% |
|  | 3.7% (n=2) | 0% | 1.4% (n=1) | Data not collected |
| Home UK | 72.2% (n=52) | 68.8% (n=53) | 62.9% (n=44) | 63.9% (n=78) |
|  | 83.3% (n=45) | 53.9% (n=41) | 74.6% (n=53) | Data not collected |
| Home EU | 9.7% (n=7) | 13% (n=10) | 11.4% (n=8) | 6.6% (n=8) |
|  | 5.6% (n=3) | 23.7% (n=18) | 7.0% (n=5) | Data not collected |
| International | 18% (n=13) | 18% (n=14) | 25.7% (n=18) | 25.4% (n=31) |
|  | 5.6% (n=3) | 9.2% (n=7) | 18.3% (n=13) | Data not collected |

Out of the five courses included, most responders were part of the BSc in Biomedical Sciences cohort, comprising 46.9-75% of the total participants (data not shown). The percentage of students responding to the study were predominately Home UK students (63.9-72.2%), followed by International (18-25.7%) and European (6.6-13%) participants (Table 1).

### B) Pre-project survey

#### 1) Student feelings towards research project

Overall, students showed mixed emotions regarding their upcoming project across the four years, with fluctuating levels of concern, dread, and apprehension, alongside positive anticipation (Figure 2). The proportion of students who expressed concern ranged 7.8-18.6%, with the 2020-21 cohort expressing the highest level of concern—a statistically significantly difference when compared to the 2021-22 cohort, which reported the lowest level of concern (P<0.05). Interestingly, a greater percentage of students indicated that they were dreading their project in 2019-20 compared to later years (P<0.05 and P<0.01, respectively). Apprehension was greatest in the 2021-22 cohort (37.7%), while the lowest level was reported in 2020-21 (P<0.001). The 2021-22 cohort expressed the lowest response rate of lack of sufficient information (1.4%) which was significantly lower compared to all other years.

**Figure 2:**
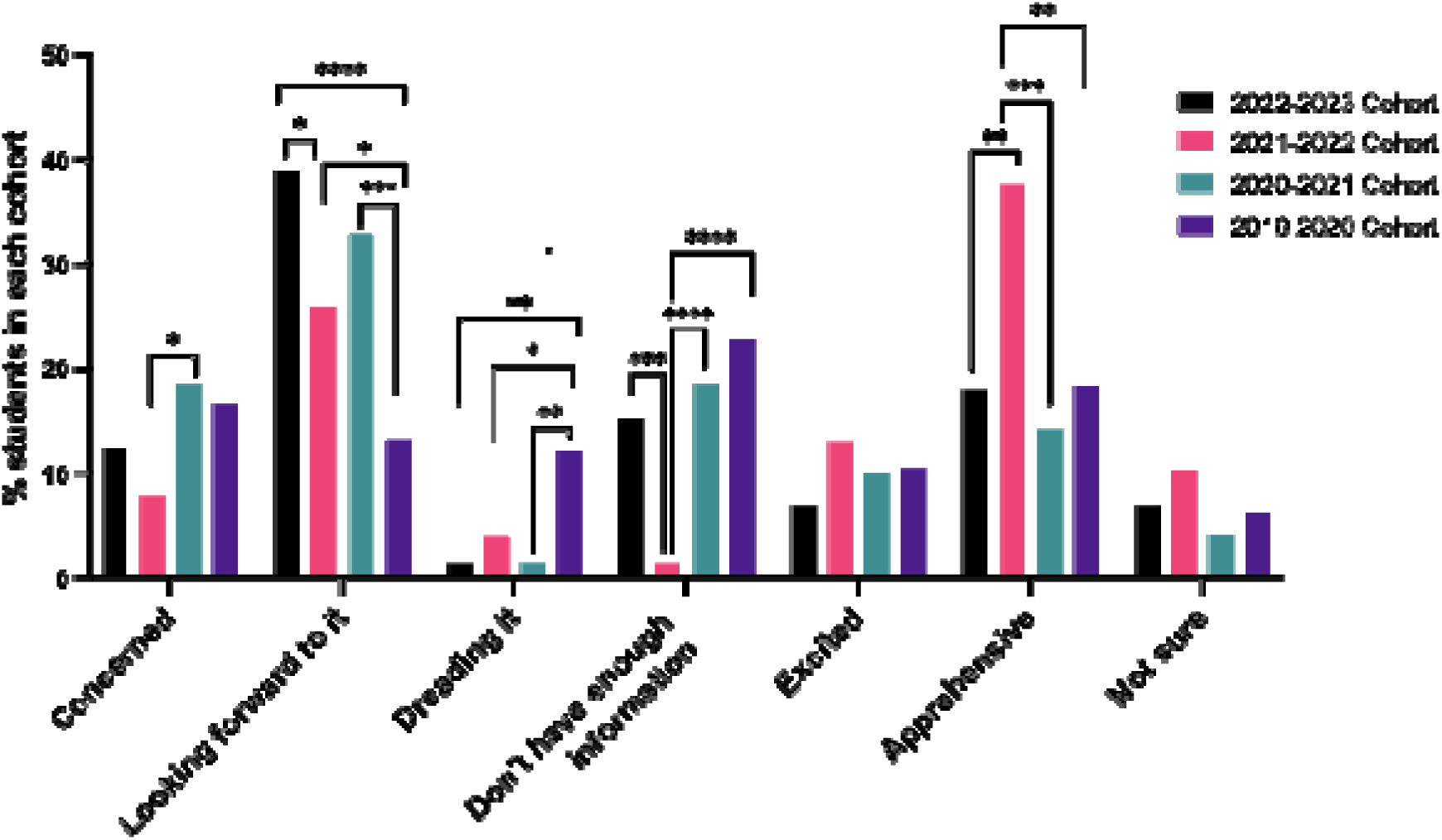
Student feelings towards research project, pre-project, 2019-2023. Statistical significance was assessed using chi squared test: P<0.0001 ****, P<0.001 ***, P<0.01 ** and P<0.05 *.

#### 2) Impact on subject knowledge

Across all cohorts, most students answered positively when asked whether they expected to advance their specific-subject understanding (80-96.6%, Figure 3). A significant decrease in 2020-21 was indicated when student responses were compared to the other three cohorts (P<0.001 and P<0.01 respectively, Figure 3). A small proportion of students across all years selected ‘no’ (0-4.3%), ‘don’t know’ (0-5.7%), or ‘unsure’ (0-10%), with the highest levels of uncertainty reported in 2020-21 (10%). In summary, the 2020-21 cohort exhibited the most negative response pattern, indicating a greater degree of uncertainty and reduced relevance of the project to their subject understanding.

**Figure 3:**
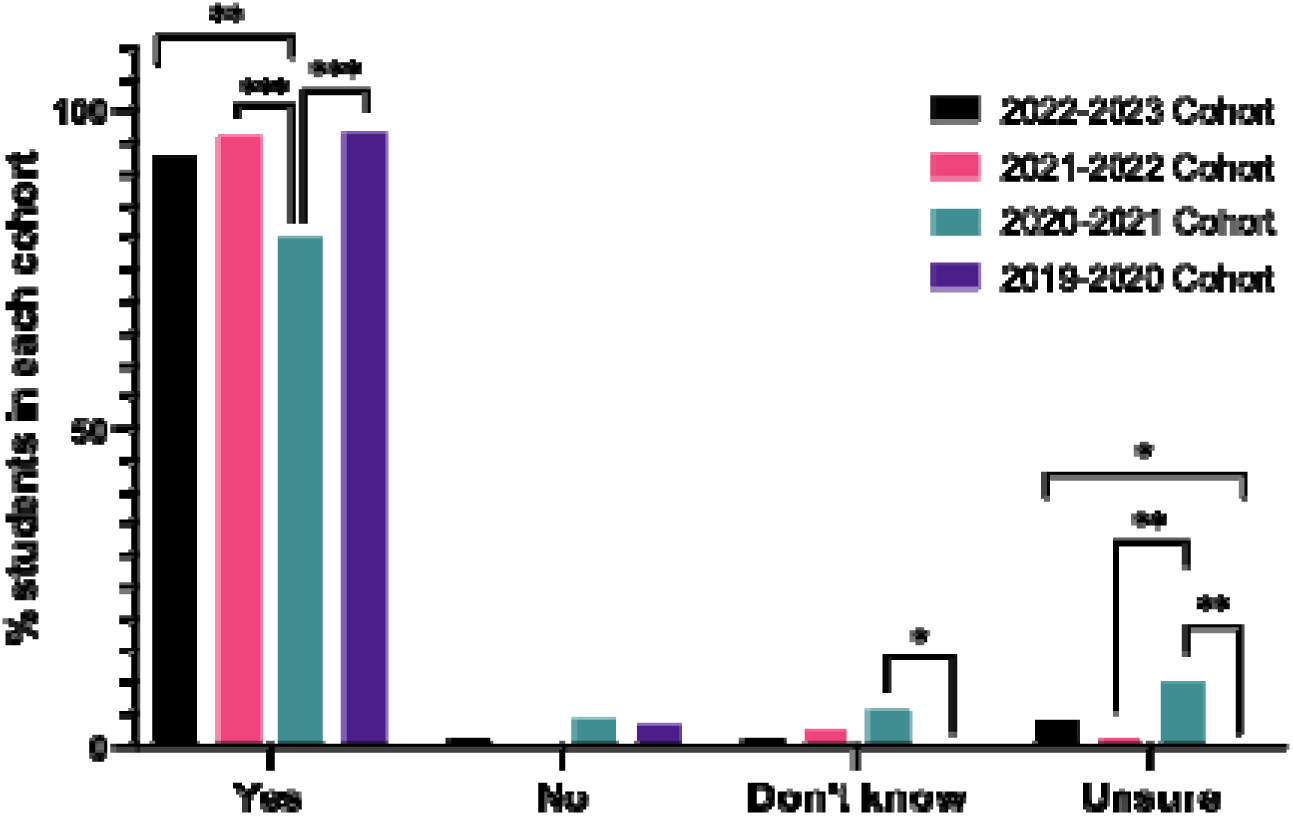
Student perceptions of the potential impact of research project on subject knowledge, pre-project, 2019-2023. Statistical significance was assessed using chi squared test: P<0.0001 ****, P<0.001 ***, P<0.01 ** and P<0.05 *.

#### 3) Perceptions on graduate attribute development

When participants were asked about the graduate attributes expected to develop during their projects, critical thinking scored highest across all years, ranging from 89.5% to 92.8% (Figure 4). On average, more than two-thirds of the students expected to develop confidence (56.6-81%). However, this was significantly lower in the 2020-21 cohort compared to other years (P<0.001). Students from the 2020-21 and 2022-23 cohorts felt less likely to develop resilience from their projects compared to students from the other two years (P<0.05). The 2020-21 cohort felt that engagement and curiosity were to be developed compared the other three cohorts (p<0.001 compared to 2022-23 cohorts). Opportunity for collaboration was reported at similar levels across all four cohorts, ranging from 63% to 67%.

**Figure 4:**
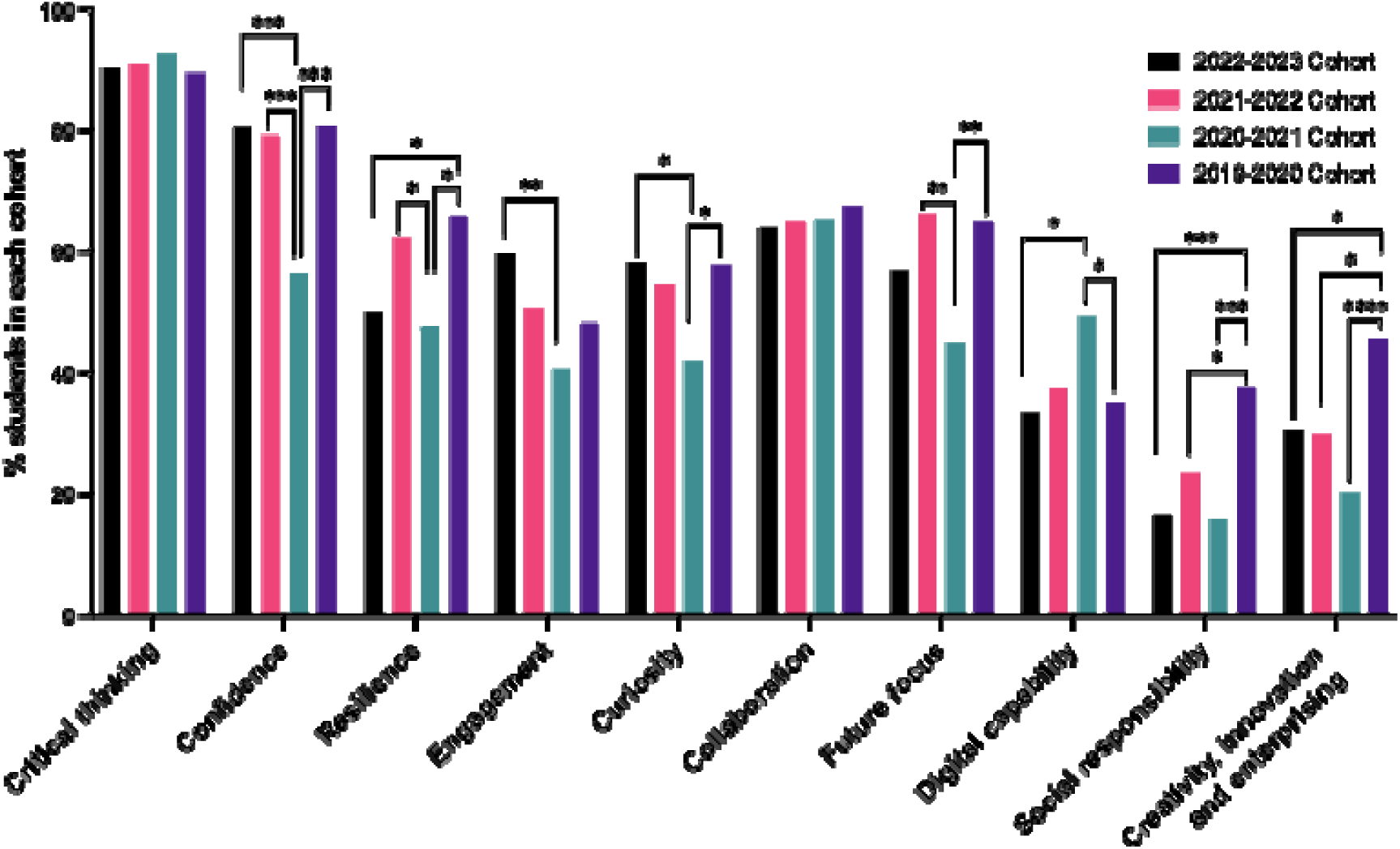
Student perceptions of graduate attribute development from their final year project, pre-project, 2019-2023. Statistical significance was assessed using chi squared test: P<0.0001 ****, P<0.001 ***, P<0.01 ** and P<0.05 *.

Our results also revealed that students from the 2020-21 cohort were thinking differently about future focus, showing significant differences from the previous and following years (P<0.01). More than one-third of the students in 2019-20 (37.7%) expected to develop social responsibility skills, an expectation that reduced significantly in the following year (15.9%) (P<0.001) and continued to decline in subsequent years (P<0.001 and P<0.05, respectively) (Figure 4).

In addition to social responsibility, the anticipated development of digital capability, creativity, innovation and enterprising were generally low ranked among students. On average, about a third of students expected to advance these skills during their dissertations. In the 2020-21 cohort, a higher proportion of students (49%) indicated they expected to develop digital capability compared to students from the other years (P<0.05, compared to both 2019-20 and 2022-23). Meanwhile, a significant increase in the percentage of students from the 2019-20 cohort expected to develop creativity, innovation and enterprising through their projects (45.6%, P<0.0001 compared to 2020-21 and P<0.05 compared to the other two years) (Figure 4).

#### 4) Future employability

Most participants expected their project to enhance their employability (62-80%). The highest percentage was recorded in the 2021-22 cohort, which displayed a statistically significant difference compared to the previous two cohorts (P<0.01). This was followed by the 2022-23 cohort, which showed a higher proportion of students answering positively compared to 2019-20 and 2020-21 (P<0.05) (Figure 5).

**Figure 5:**
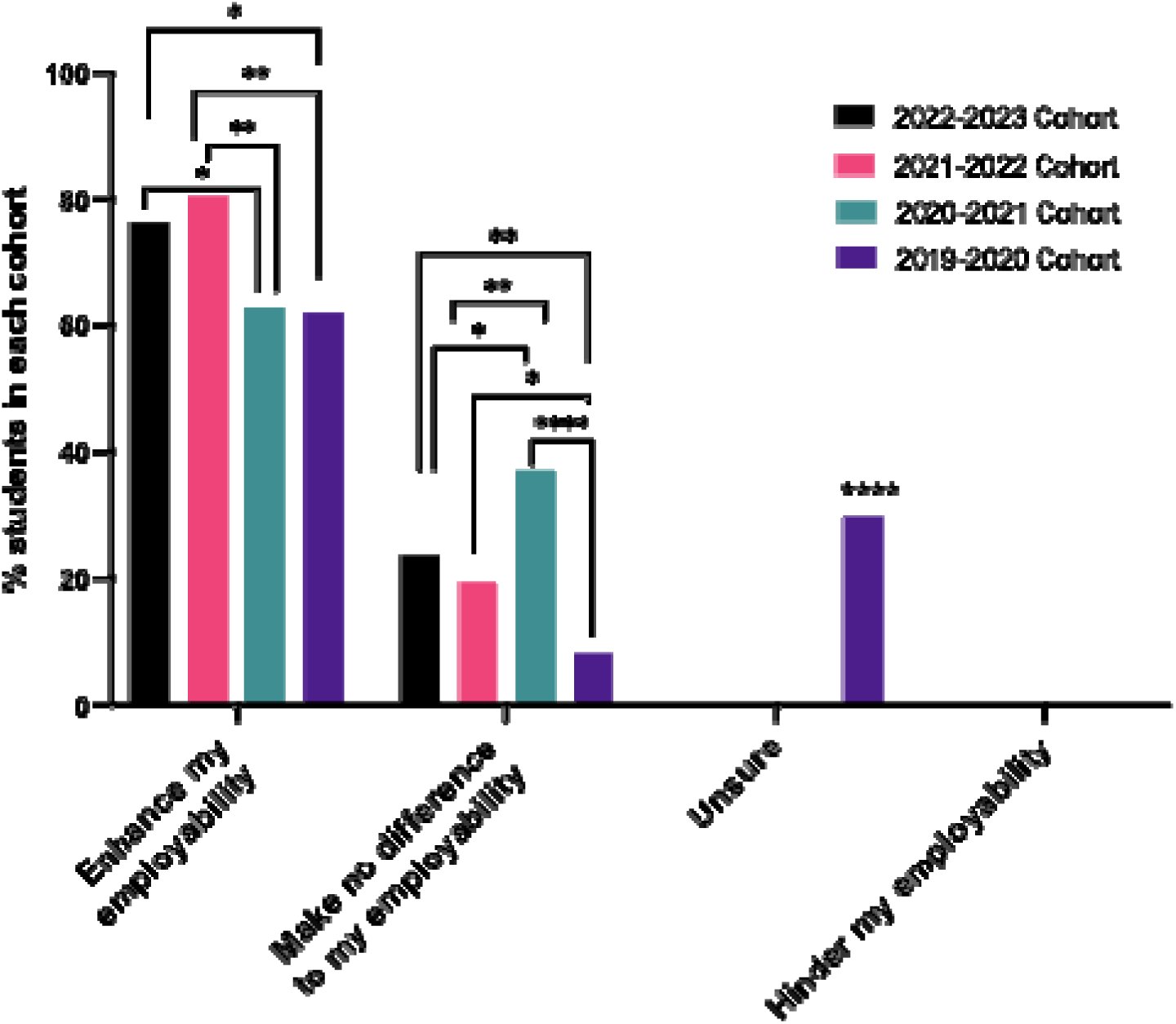
Student’s perceptions of potential impact of project on their future employability, pre-project, 2019-2023. Statistical significance was assessed using chi squared test: P<0.0001 ****, P<0.001 ***, P<0.01 ** and P<0.05 *. For unsure, P<0.0001 when 2019-20 compared to other 3 years. No students across all four cohorts thought the project would hinder their employability.

The lowest percentage of responders who expressed that the project would make no difference to their employability was recorded in 2019-20 (8%), while the highest was from the 2020-21 cohort (37.1%) (P<0.0001 when compared). The percentage responding from the 2020-21 cohort was significantly higher compared to all three other cohorts (P<0.05 to P<0.00001, respectively). Among the 2019-20 cohort, one third of the students (30%) were unsure about the project’s impact on employability, which was significantly increased compared to the other cohorts, where no students expressed this uncertainty (P<0.0001) (Figure 5). Across all four cohorts, none of the students expected their project to hinder their employability.

### C) Post-project survey

#### 5) Student feelings towards research project

Most students reported enjoying their project, with student enjoyment reaching 69.5% in 2019-20, and rising to 88.9% in 2022-23 (data not shown). Between 7.3% and 16.9% of students reported lack of enjoyment, with the peak of negative response emerging from the 2020-21 cohort. A small percentage of students remained uncertain about their project experience (0-2.6%), with the highest proportion of those coming from the 2020-21 cohort and ranging between 3.7 - 11.3%.

When questioned whether the research project met their expectations, most students responded positively (50.7-77.8%). However, the 2020-21 cohort expressed a significantly lower level of satisfaction compared to other cohorts (P<0.0001 to P<0.05, respectively, Figure 6). A greater percentage of students from 2020-21 also indicated that their project did not meet their expectations (28.2%), which was significantly higher compared to the other cohorts (P<0.0001 to P<0.05, respectively). Only a small proportion of students answered, ‘don’t know’ (1.9-6.6%) across all cohorts, but the levels of uncertainty varied, with the 2021-22 group reporting the lowest levels compared to the subsequent and previous years (P<0.01).

**Figure 6:**
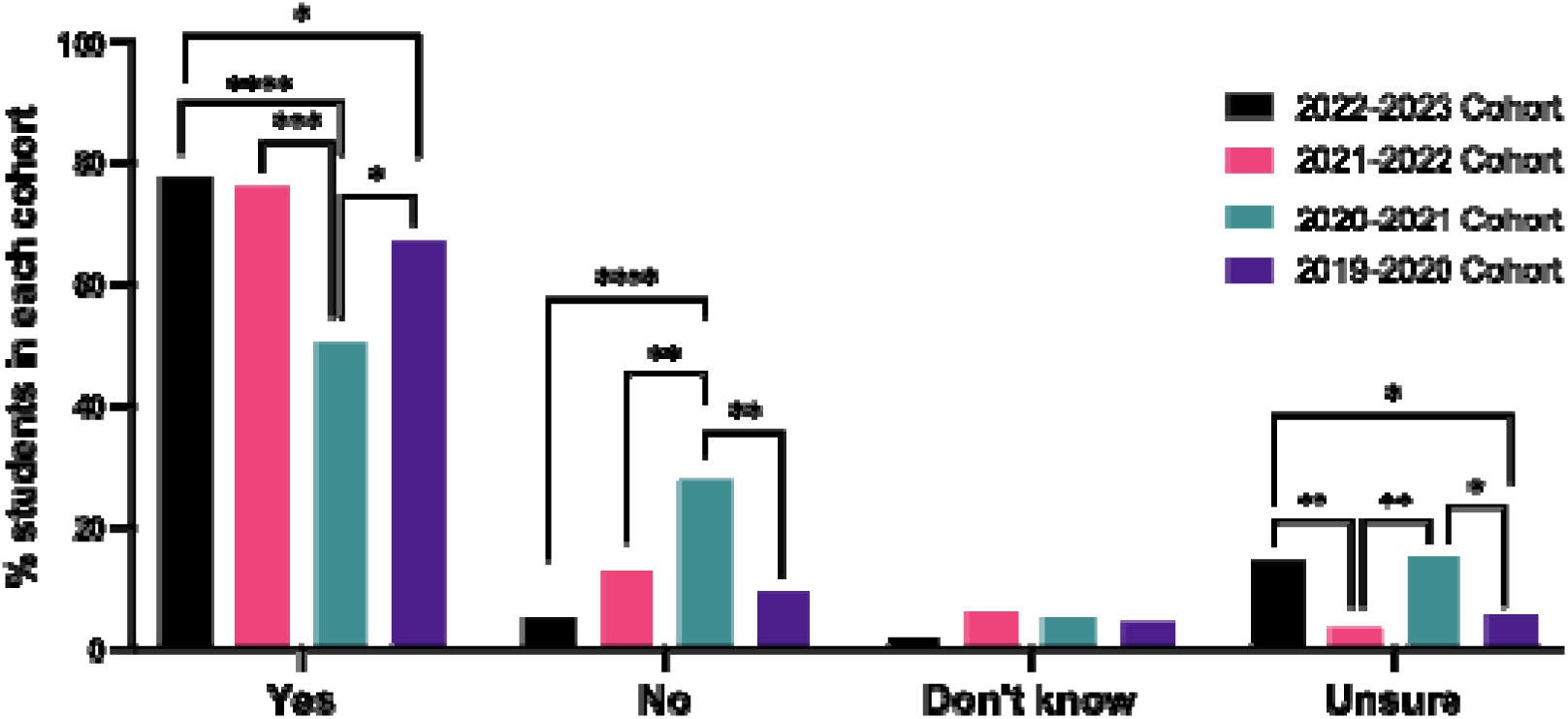
Expectations of project met, post-project, 2019-2023. Statistical significance was assessed using chi squared test: P<0.0001 ****, P<0.001 ***, P<0.01 ** and P<0.05 *.

#### 6) Subject knowledge impact

When asked whether subject knowledge would be enhanced by the research project, similar mean averages across cohorts were reported, with the majority answering positively. Specifically, 91.4% of students indicated their project to have an impact on the subject knowledge before their project started, a percentage that increase to 94.4% post-project (data not shown).

#### 7) Perceptions on graduate attribute development

We next sought to determine the graduate attributes students developed following project completion; critical thinking, confidence and resilience were most selected (Figure 7). Fewer students in 2019-20 selected resilience as a graduate attribute compared to other cohorts (P<0.01 compared to 2020-21 and P<0.05 to 2022-23).

**Figure 7:**
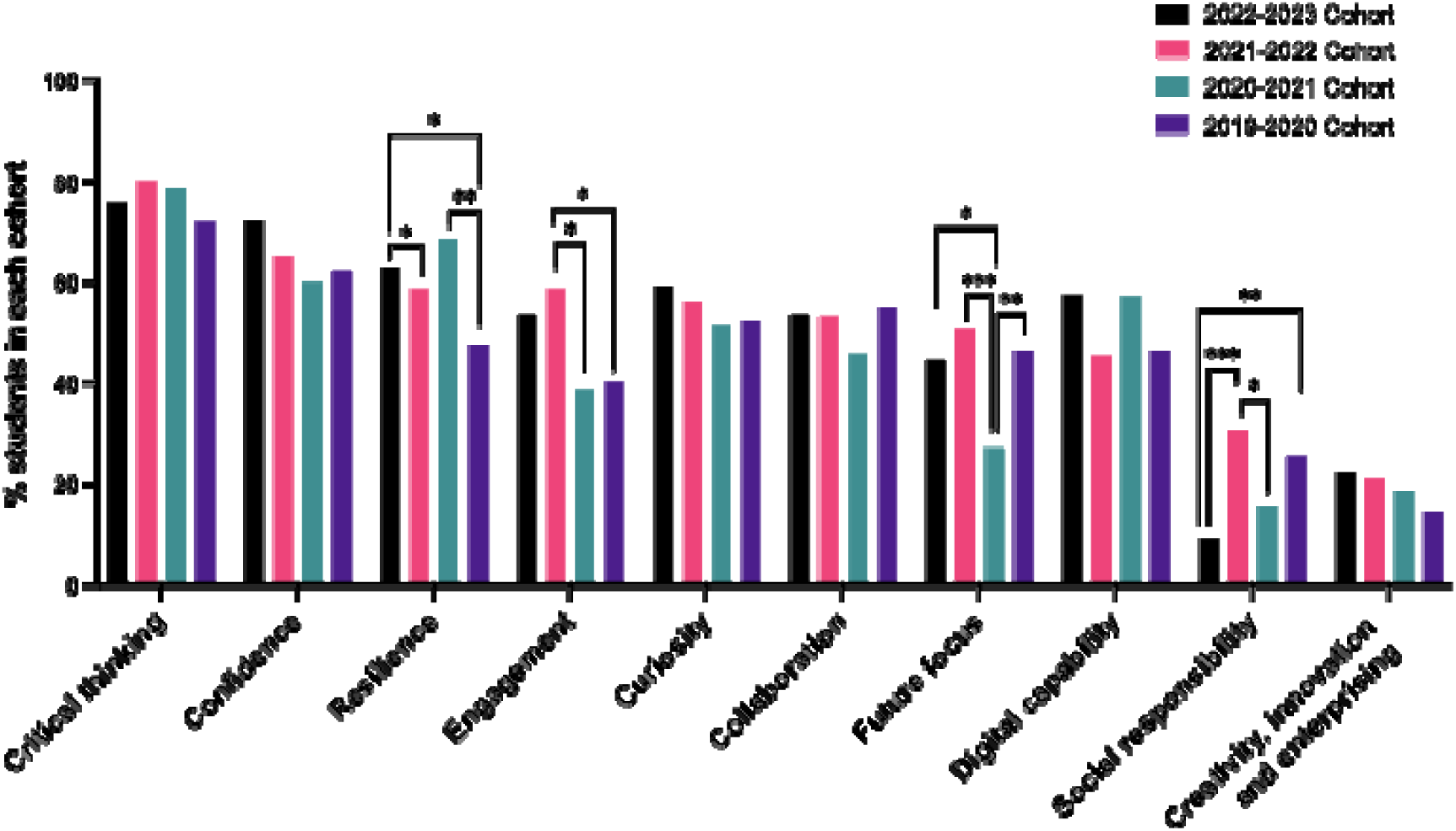
Student’s perceptions of graduate attribute development from their final year project, post-project 2019-2023. Statistical significance was assessed using chi squared test: P<0.0001 ****, P<0.001 ***, P<0.01 ** and P<0.05 *.

Notable variabilities were observed on engagement as a graduate attribute developed with the lowest levels recorded in the 2019-20 and 2020-21 cohorts (P<0.05 when compared to 2021-22). Similar response rates for curiosity and collaboration were recorded across the cohorts. However, future focus was significantly lower in the 2020-21 cohort compared to other years (P<0.001 to P<0.05, respectively).

Across cohorts, almost a half of students indicated they had developed digital capability through their project (45.3-57.4%). In contrast, perceptions on social responsibility were highly variable with the lowest levels reported in 2022-23 (9.3%), and 2020-21 (15.7%) (P<0.05, P<0.01 and P<0.0001 compared to other years) and highest in 2021-22 (30.7%).

#### 8) Graduate attributes pre and post project

A comparison between students anticipated (pre-project) and actual (post-project) GAs developed were compared (Figure 8) and revealed broadly similar profiles across most skills. However, notable differences were recorded for specific attributes. Critical thinking development post-project was significantly lower than pre-project (75.9% compared to 90.3%, P<0.01). A similar trend was also noted in future focus, which was significantly lower upon project completion (P<0.001). In contrast, digital capability showed the opposite trend, with 57.4% of students recognising development of this skill after project completion, compared to 33.3% pre project (P<0.001).

**Figure 8:**
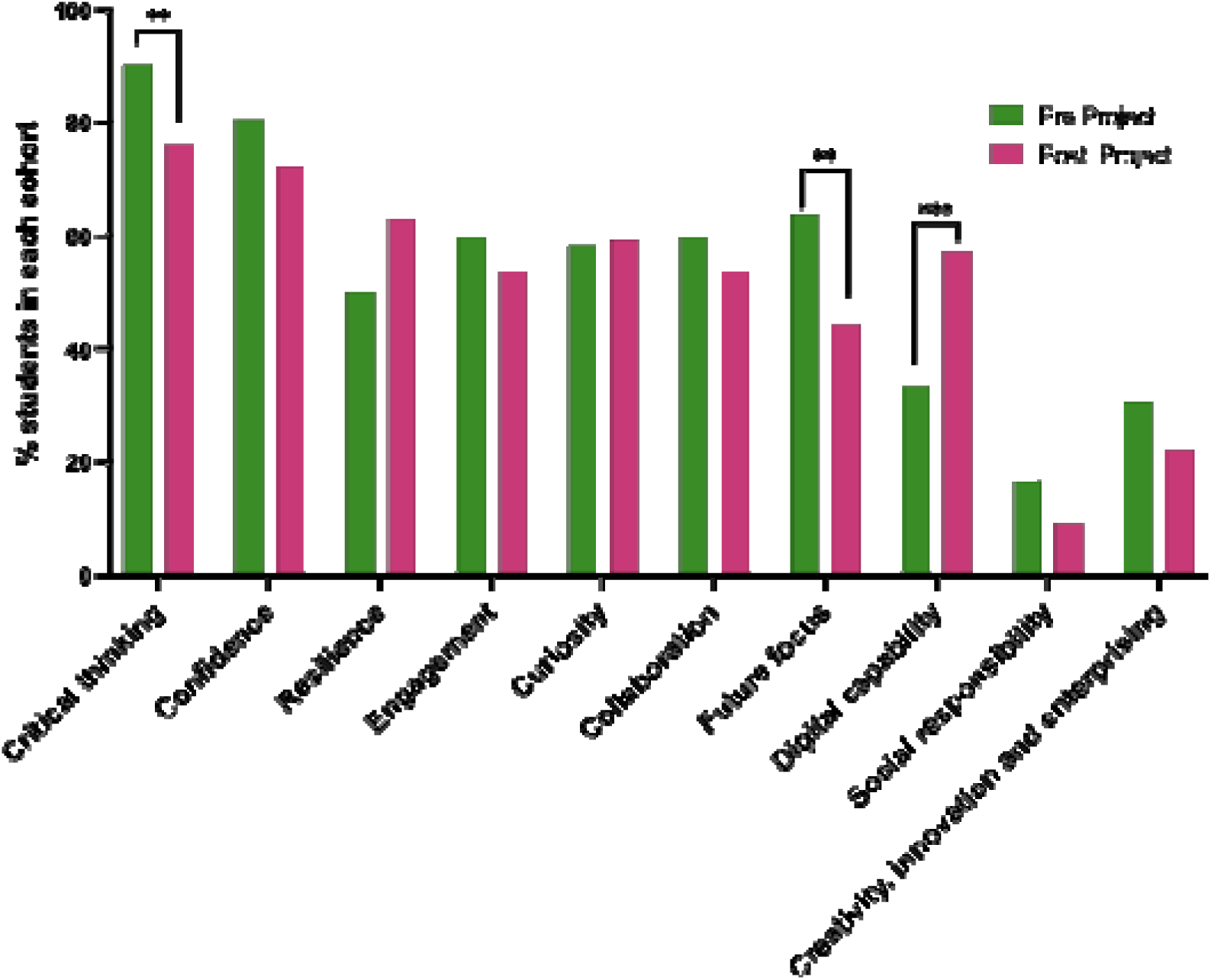
Student’s perceptions of graduate attribute development from their final year project, pre- and post-project 2019-2023. Statistical significance was assessed using chi squared test: P<0.0001 ****, P<0.001 ***, P<0.01 ** and P<0.05 *.

#### 9) Future employability

When we assessed employability impact, approximately three quarters of students perceived that the project enhanced their employability (70.0-77.8%). A smaller proportion (20.3-28.6%) indicated that the project made no difference to their employability and 0-1.9% expressed that the project had hindered their employability.

#### 10) Pre- and post-project employability

When perceptions on employability pre and post project were compared, similar response rates were recorded. Prior to project commencing, 70.4% of participants expected their project to enhance their employability increasing to 75.0% after the project completion. Similarly, the percentage of those expressing that their project would make no difference to their employability remained relatively stable, with 22% pre-project and 24% post-project. These findings suggest that student expectations on employability were largely met upon project completion.

## Discussion

Undergraduate students are expected to develop a broad range of skills and experiences to support them academically and promote their post-graduation employability, particularly in securing graduate level roles. Undertaking a final year research project provides an authentic enquiry-led assessment which can be classed as high stakes to undergraduates and contributes a significant number of credits for final year and degree classification. We demonstrate that a high proportion of students enjoy their research project (69.5-88.9%), showing an upward trend over time. Whilst this study focused on GAs, it is equally important to note that these are not independent of the development of research competencies or subject-specific knowledge in the biosciences. We provide evidence demonstrating that most students acknowledge the potential of the project in developing their subject knowledge (91.4% and 94.4% pre- and post-project respectively, Figure 3).

Providing sufficient capstone projects can be particularly challenging due to the increasing size of student cohorts and funding opportunities within universities. This is particularly pertinent within life sciences where traditionally projects have been within a laboratory setting (Linn et al., 2015; Wannapiroon, 2014). However, as the field has developed and to support the diversity of students’ interests and aspirations, a wider range of project types have been introduced, for example, computational and large data set analysis. Importantly, standardising the learning experience across a broad range of project types and topics, increasing student numbers and with many staff involved, can be challenging for institutions.

The COVID-19 pandemic had a huge impact on education and teaching delivery. From March 2020 onwards, there was a rapid unplanned shift to remote teaching and assessment, accompanied by a high level of uncertainty (Kerr et al., 2023; Rauseo et al., 2022; Sustarsic & Zhang, 2022). With regards to the research project, this meant a shift to remote projects with very limited time to prepare for or to predict the impact of this change. All project introductions, support meetings, feedback were delivered remotely for 2020-21. This, therefore, constituted a very different learning experience altering not only the type of projects provided, but also how students engaged with their supervisors and research teams.

The 2020-21 cohort had the most negative perception on project contribution and subject knowledge enhancement (Figure 3). Notably, they were less likely to recognise resilience as a skill developed compared to other cohorts but acknowledged and anticipated an increase in their digital capabilities (Figure 4). This group also reported the lowest perceived impact of the project on employability (Figure 5) and post-project, the highest level of dissatisfaction of the project experience. Although in person delivery of projects was restored in 2021-22, the levels of uncertainty derived from the pandemic, remained in the first semester, likely contributing to the high levels of apprehension recorded in the 2021-22 cohort (37.7%, Figure 2). It is also important to note that teaching delivery across the whole of the programme was impacted March 2020 to September 2021, this involved switching from in person sessions to online delivery including laboratory sessions. This may therefore have negatively impacted students’ practical skill confidence and preparedness of their research project (Grineski et al., 2022; Watts et al., 2022).

Previous literature has evidenced mixed findings regarding skills development via research projects across a range of disciplines, often with disparities between student and supervisors’ perceptions of what has been developed (Brewer et al., 2012; Yeoman & Zamorski, 2008). However, alumni surveys highlight the value of dissertations, and the skills developed during this period of research. Therefore, it may be that students do not have the time or perhaps tools, to adequately reflect and recognise their own skill development during the time of their studies (Bauer & Bennett, 2003).

Critical thinking was a key attribute that large proportions of students both expected to develop and reported developing through their research project (Figures 4, 7 and 8) although it appears that their expectations of this skill development wasn’t fully achieved (90.3% pre-project compared to 75.9% post-project). For the other nine GA’s, a notable variation was observed between cohorts. Most of the lower response rates were reported by the 2020-21 cohort (confidence, resilience, future focus engagement, curiosity, creativity) likely reflecting lower expectations because of COVID-19 and the change to remote projects. However, this cohort reported higher anticipated development of digital capabilities (49.3%), perhaps recognising the project and platform use changes. Previous literature suggests that individuals with a high level of digital skills were able to maintain focus and engagement during the COVID-19 lockdowns and performed better academically (psychology and veterinary science) (Limniou et al., 2021).

Post-project GA trends were comparable to pre-project, but interestingly the 2020-21 cohort reported higher resilience development post-project compared to the other cohorts; again, perhaps related to the impact of the pandemic (Figure 8). When comparing the mean scores across all four academic cohorts, critical thinking, future focus and digital capability were significantly different between pre- and post-project responses (Figure 8). However, for critical thinking and future focus, the post-project response levels were lower than the pre-project expectations, suggesting that the project experience didn’t meet students’ expectations for these two GAs. Notably, the increase in recognition of digital capability was a positive outcome considering the substantial use of digital tool and platforms throughout the project (i.e., 5000-word report, data analysis, and oral presentation). Digital skills remain a key area of focus in HE with constant updates and developments. The economic and professional consequences of insufficient digital skills have been reported and account for the loss of over £23 billion per year in UK (Future.now, 2025). Moreover, the reported graduate and employers’ skills gap also highlights the need for focusing on digital skill development (Commision, 2017; Education, 2023).

Regarding future employability, the majority of students felt that the project would have a positive impact pre-project (Figure 5), but there was a significant difference between 2019-20 and 2020-21 compared to 2021-22 (62% and 62.9% for earlier years compared to 80.5% in 2021-22 and 76.4% in 2022-23). A significantly higher percentage in 2020-21 also reported that the project would make no difference to their employability, which could again be explained by the enforced changes due to COVID-19. This finding is supported by studies that investigated student perception on their employability as a result of the pandemic (Capone et al., 2021; Ren et al., 2024). Interestingly, nearly a third of students in 2019-20 were unsure on whether the project would impact their employability. Post-project, around three quarters of students felt that their employability had been enhanced by the project, and on average there was a small increase from 70.4% to 75% following project completion. This finding doesn’t directly correlate with GA development reported earlier and perhaps suggest that there may be a disconnect in GAs, employability and the final year research project. Again, it was reassuring to see a positive acknowledgment of the research project impacting employability.

The integration of GAs into the curriculum is essential to ensure that all students are aware of their definition and how they can develop them (HE, 2024). Also, aligning GAs directly to employability, ensuring what these skills are and supporting students in articulating them is fundamental to effective implementation in student outcomes. Clear and consistent terminology needs to be established from the outset of student programmes to avoid ambiguity and to ensure consistency is maintained across all stakeholders (HE, 2024). Although not addressed within this project, the authors have concerns that not all students fully understand the definition of GAs and how they have developed them, particularly in reference to their research project. How these GAs are captured, reflected upon, and achieved by graduation remains an ongoing topic of discussion. Newcastle University has recently reviewed its GAs, now identifying 15 attributes/skills with significant cross-over with the 10 reported in this study. Importantly, the provided surveys created an opportunity for students to consider and reflect upon their GAs and think about their upcoming research project and personal growth.

## Limitations

Response rates to the two surveys fluctuated across the years (16.2%-35.9%), with the greatest proportion of responses being via an in-person session. Due to logistical and timing constraints, this was not repeated. Demographics of the respondents was generally representative of the cohort but there was an overrepresentation of females (average 67.6% female in cohorts over period of study) and those who identified as non-UK (internal data). As some cohorts had low rates, and the surveys were not compulsory, there is the potential that the data is not truly representative of the cohorts but instead a self-selecting group. Falling student response rates has also been reported in the literature, possibly due to ‘survey fatigue’ but also that these surveys were run at busy academic periods within the RQF level 6 year (Nair et al., 2008).

Both surveys used self-assessment of graduate attributes as opposed to teacher assessed ratings with no standards of what the expected level of each skill is, bias could therefore be a factor (Butler, 2011). Gathering supervisor feedback on their student’s skills would be an interesting angle to take alongside evidence of the reported skills, for example aligning skills with project marks. As both surveys were carried out anonymously and no identifiers were given to each respondent, it is not known whether those who completed the pre-project survey were the same or similar group within the cohort. The surveys also presumed that students had a full understanding of the terminology of GAs, but since this was not investigated fully, students were only signposted to the University’s Graduate Framework site for reference (University, 2018).

## Conclusions

Our data demonstrate that the final year research project does create concerns. Consequently, over the years additional support sessions that are scheduled earlier in semester 1 of Level 6 have been introduced, ensuring that students are better prepared for their upcoming project. School-based student well-being advisers have also since been introduced to the School and a range of sessions and targeted support for finalists during their research project have been well received. Consistency of experience is very difficult to achieve with large cohorts alongside variation in project types, and so this may explain some of the research findings. This was also commented upon in many of the free text comments (data not included). Clearly students recognise a wide range of skills that they develop via their project and generally find the experience to be positive. The COVID-19 pandemic showed a significant impact on student experience, confirmed by our findings, creating high levels of uncertainty and negativity across many groups. The final year project provides students with a unique opportunity to pose a research question and develop a range of skills but whether these are fully recognised by students needs further exploration.

## Abbreviations

GAs: Graduate Attributes
HEI: Higher Education Institute
COVID-19: SARs-Cov2 pandemic
QAA: Quality Assurance Agency

## Data accessibility

Data will be made available on request to the corresponding author.

## Author contributions

VA, JJ and SW conceived the study; VA supervised and co-ordinated the study; MK, VA and KP analysed data, SW provided statistical advice; VA and MK wrote the manuscript, VA, MK, JJ, KP, SW and AB supported manuscript revisions.

## Acknowledgements

We are grateful to have received funding from Newcastle University’s Education Development Fund to initiate this project. Thanks to previous project interns Ramandeep Dhanoa, Emily Jeffreys and Rebecca-Leigh Railton.

